# Mathematical diabetes disease progression modeling in the integrated glucose-insulin model among individuals with impaired glucose tolerance from the Finnish Diabetes Prevention Study

**DOI:** 10.64898/2026.02.28.708712

**Authors:** Siti M. Sheikh Ghadzi, Mats O. Karlsson, Vanessa D. de Mello, Matti Uusitupa, Maria C. Kjellsson, Finnish Diabetes Prevention Study Group

## Abstract

The integrated glucose-insulin (IGI) model describes glucose and insulin after glucose administration in healthy individuals and patients with type 2 diabetes. The model, however, does not include disease progression (DP) from prediabetes to overt diabetes, which is driven by decreased insulin sensitivity and relative beta-cell failure. The objective of this study was to develop the IGI model to include the DP model for glucose and insulin in individuals with impaired glucose tolerance (IGT), with and without lifestyle intervention. Data of frequently sampled intravenous glucose tolerance test and oral glucose tolerance test (OGTT) were obtained from a sub study of the Finnish Diabetes Prevention Study (FDPS) in 101 individuals with IGT, randomly assigned to control and lifestyle intervention groups. A combination of intravenous and oral IGI model was used to fit the baseline until the fourth-year data using NONMEM, with prior information. The first-phase insulin secretion (FPS) and insulin-dependent glucose clearance (CL_GI_) decreased by 3.0% year and 8.1%/year, due to DP. Baseline insulin concentration (I_SS_) was increased by 68% from baseline to Year 1, and remained unchanged thereafter. With intervention, a net reduction of 0.1%/year for FPS and reduction of 2.1%/year for CL_GI_ was quantified, translated to a much slower deterioration of the first-phase insulin secretion and insulin sensitivity. The I_SS_ was affected by a net increase of 153% from baseline to Year 1 and remained constant after that, possibly reflecting beta-cell function improvement. The DP was successfully included in the IGI model to describe differences in IGT population, with and without lifestyle intervention.

## Introduction

The integrated glucose-insulin (IGI) model was published^1–3^ to describe the glucose and insulin regulation system in various glucose provocations, e.g. intravenous (IVGTT) and oral glucose tolerance tests (OGTT) among healthy individuals and also patients with type 2 diabetes mellitus (T2DM). A similar model structure is used to describe the IVGTT and OGTT, with differences in the insulin-independent glucose clearance, and with the addition of incretin effect and glucose bioavailability in the OGTT model. In the healthy and diabetic populations, except for insulin-dependent and insulin-independent glucose clearance (CL_GI_ and CL_G_, respectively), first-phase insulin secretion (FPS), rate constant of FPS (k_IS_), the feedback mechanism of glucose effect on its own production (GPRG), incretin effects, and glucose bioavailability (BIO_G_).^1–3^ Although being able to describe both healthy volunteers and diabetes patient populations, the disease progression (DP) from healthy to diabetic which is driven by decreased insulin sensitivity and relative beta-cell failure as in individuals with impaired glucose tolerance (IGT), is not included in the model. The inclusion of a DP model then is deemed important to quantify those changes over time.

IGT is a pre-diabetic state that exists as a result of abnormal glucose homeostasis, in between the states of normal glucose tolerance (NGT) and diabetes.^4^ Based on the new World Health Organization (WHO) Global Report on Diabetes^5^, IGT is defined as a fasting plasma glucose (FPG) of <126 mg/dL and 2-hour plasma glucose between ≥140 mg/dL and 200 mg/dL after a 75 g OGTT. Another form of pre-diabetic state is known as impaired fasting glucose (IFG), defined as FPG concentration from 110 mg/dL to 125 mg/dL and (if measured) 2-hour plasma glucose concentration of <140 mg/dL.^5^ Both IGT and IFG are in insulin-resistance states, but people with IGT have a moderate to severe muscle insulin resistance and slightly reduced hepatic insulin sensitivity, whereas people with IFG have a normal muscle insulin resistant but a high hepatic insulin resistant.^4^ Results of studies comparing the two pre-diabetic states are inconclusive regarding the differences between changes insulin resistance, first and second phase insulin secretion related to the beta-cell function, and incretin function between IGT and IFG.^6–12^ In this current study, however, the focus of the discussion will be on individuals with IGT, as the studied population.

Diabetes DP is clinically characterized into progression from pre-diabetes to overt diabetes, by the lack of acute insulin response, declining of beta-cell function measured by clinical tools for example homeostasis model assessment (HOMA), progression to medication, loss of glycemic control on medication, as well as progressive weight gain.^13^ Based on the glucose level measurement, the natural history of pre-diabetes can be explained by the development of progressive hyperglycemia, which eventually meets the criteria for diagnosis, as seen in the majority of people in the pre-diabetes population. The hyperglycemia condition is due to the progressive decline in insulin secretion or increase in insulin resistance among pre-diabetic individuals.^4^ Failure of acute insulin response occurs when the insulin level has failed to be increased by the compensatory mechanism, which then leads to IGT and a decreased in insulin secretion, and later, diabetes.^14^ The declining of the beta-cell function is a major sign of diabetes disease progression, which begins around 12 years before diagnosis.^13^

The beneficial effect of lifestyle intervention such as changes of dietary intake (increasing fiber, and reducing fat intake), weight reduction management, as well as physical activities (such as moderate intensity exercise for at least 30 minutes, 5 times a week) has previously been investigated. An improvement in glucose tolerance, beta-cell function and insulin sensitivity was documented with the lifestyle intervention, due to weight loss, and increasing physical fitness.^15–17^ A significant reduction of the progression from IGT to T2DM was also reported in the lifestyle intervention group throughout the mean follow-up duration of 2.6 and 3.2 years.^18,19^

With regard to the importance of quantifying the physiological changes of the progression from pre-diabetes to overt diabetes, this study was done to develop the IGI model to include DP model on the glucose and insulin components among individuals with IGT. The developed model was used to quantify the effect of natural DP and lifestyle intervention on the parameters related to the changes in glucose and insulin regulations.

## Methods

### Data

Data from a sub study of the Finnish Diabetes Prevention Study (FDPS),^19,20^ involving 101 middle-aged (with mean age of 53 years old) and overweight (with mean Body Mass Index (BMI) of 31.5) individuals with IGT were used for analysis. IGT was defined as a plasma glucose concentration of <140 mg/dL (<7.8 mmol/dL) after an overnight fast, and 140 to 200 mg/dL (7.8 to 11.0 mmol/L) 2 hours after the oral administration of 75 g of glucose.^19,21^ The individuals were randomly assigned into two groups, which were control and lifestyle intervention groups. The individuals in the control group were given general oral and written (a two-page leaflet) information about diet and exercise at baseline and annual visits, without any specific individual program. On the other hand, individuals in the intervention group were given intensive individual lifestyle interventions with counselling on weight reduction, choices of diet, and increasing physical activity with associated goals of interventions. The goals of interventions were 5% reduction of weight, total intake of fat and saturated fat less than 30% and 10% of energy consumed, increased in fiber intake to at least 15g per 1000 kcal, and moderate exercise for at least 30 minutes per day.^19,20^

All individuals underwent yearly OGTT at Year 0 (baseline) until the end (Year 4) of the intervention. Individuals who developed diabetes were excluded from the study at the time of diagnosis, after a second OGTT was done for the diagnosis confirmation, based on the 1985 WHO technical report series^21^. The diabetes diagnosis was defined as either an FPG concentration of ≥140 mg/dL or 2-hour plasma glucose concentration of ≥200 mg/dL after a 75g OGTT.^21^ Besides the yearly OGTTs, 87 out of 101 individuals underwent frequently sampled IVGTT at Year 0 and 52 individuals at Year 4. Sparse blood sampling was performed for OGTT, which samples at baseline (0 min) and every 30 mins for 120 mins after the glucose dose; and intense after IVGTT, which was at baseline (-5 and 0 min) and 23 times after the glucose dose (at 2, 4, 6, 8, 10, 12, 14, 16, 19, 22, 24, 27, 30, 40, 50, 60, 70, 90, 100, 120, 140, 160, and 180 minutes). The plasma glucose was measured with a standardized method in a laboratory in Helsinki, and the serum insulin was measured by radioimmunoassay at Pharmacia, Uppsala, Sweden. A written informed consent was obtained from the individuals and the study was approved by the ethics committee of the National Public Health in Helsinki, Finland.^19,20^

### The IGI model among individuals with IGT

To describe the IGI model among individuals with IGT, the combination of intravenous and oral IGI model^1–3^ (Figure 1) was used to fit the data for baseline, incorporating prior information on the parameters from the published IVGTT and OGTT-IGI models^1–3^ using prior^22^ functionality ($PRIOR) in NONMEM (version 7.3)^23^.

**Figure 1:**
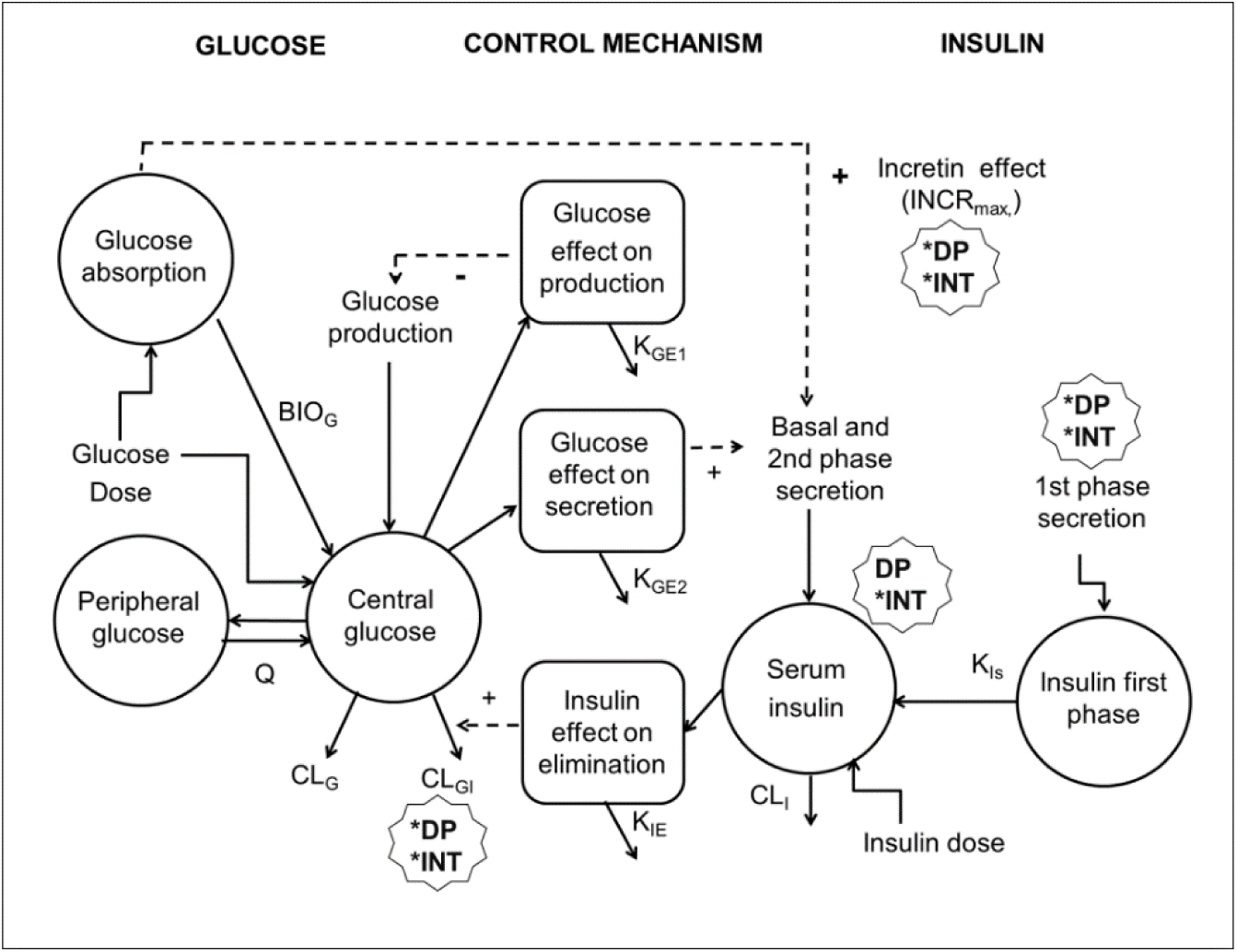
The combination of oral and intravenous integrated glucose-insulin (IGI) model^1–3^ with the disease progression (DP) and lifestyle intervention (INT) effects model on the first phase insulin secretion (FPS), insulin-dependent glucose clearance (CL_GI_), baseline insulin concentration (I_ss_), and maximum incretin effect (INCR_max_). The final model was chosen with a linear DP and INT for FPS and CL_GI_, and a step function from the baseline to the first year for I_ss_, in which the I_ss_ level was unchanged after the first year. No significant effects of DP and INT were found for the INCR_max_.

The parameters with the same values for oral, intravenous, healthy volunteers and patients with diabetes mellitus were re-estimated for the IGT population, with prior information. These values are assumed to be the best estimation of IGI model regardless of the types of glucose provocations and population. In addition, the parameters that differed in previous populations of the IGI model between provocations or populations were estimated without prior information. These parameters were the CL_G_, CL_GI_, FPS, GPRG, BIO_G_, k_IS_, maximum incretin effect (INCR_max_), absorbed glucose at 50% maximum incretin effect (ABSG_50_), as well as baseline glucose (G_SS_) and insulin (I_SS_) concentrations, with their variability.^1–3^ The estimation was done with a lower boundary of diabetics’ value and upper boundary with healthy individuals’ value, to mimic the physiological state of the IGT population, that is not as optimum as healthy state, but also not as deteriorated as the patients with diabetes.^24^ In those cases where the parameter values approached a boundary, the parameter value was fixed to the value of the boundary for stability reasons.

Besides than the typical parameter estimates and their variability, parameter correlations were also investigated in the model, which were the off-diagonal matrix between the central glucose volume of distribution (V_G_), inter-compartmental glucose clearance (Q) and insulin volume of distribution (V_I_), as the published IGI model.^1–3^ Specific to IGT population in this study, an additional off-diagonal matrix was investigated for the correlation between CL_GI_, FPS, I_SS_ and G_SS_.

All the initial parameter values and priors were summarized in Table 1, with the parameter uncertainty or the degree of freedom (df) related to every prior value. The $PRIOR functionality is a restricted maximum-likelihood function for constraining parameter estimation based on the prior information. The Normal-Inverse Wishart Prior (NWPRI) was used, involving a normal distribution for fixed effect parameters (typical parameters) and an Inverse-Wishart for the inter-individual variability (IIV).^22^ Parameter uncertainty of the fixed effects and the df for the priors of the random effects were calculated using the reported values of relative standard error and parameter estimates from the publications of the IGI model.^1–3^

**Table 1.**
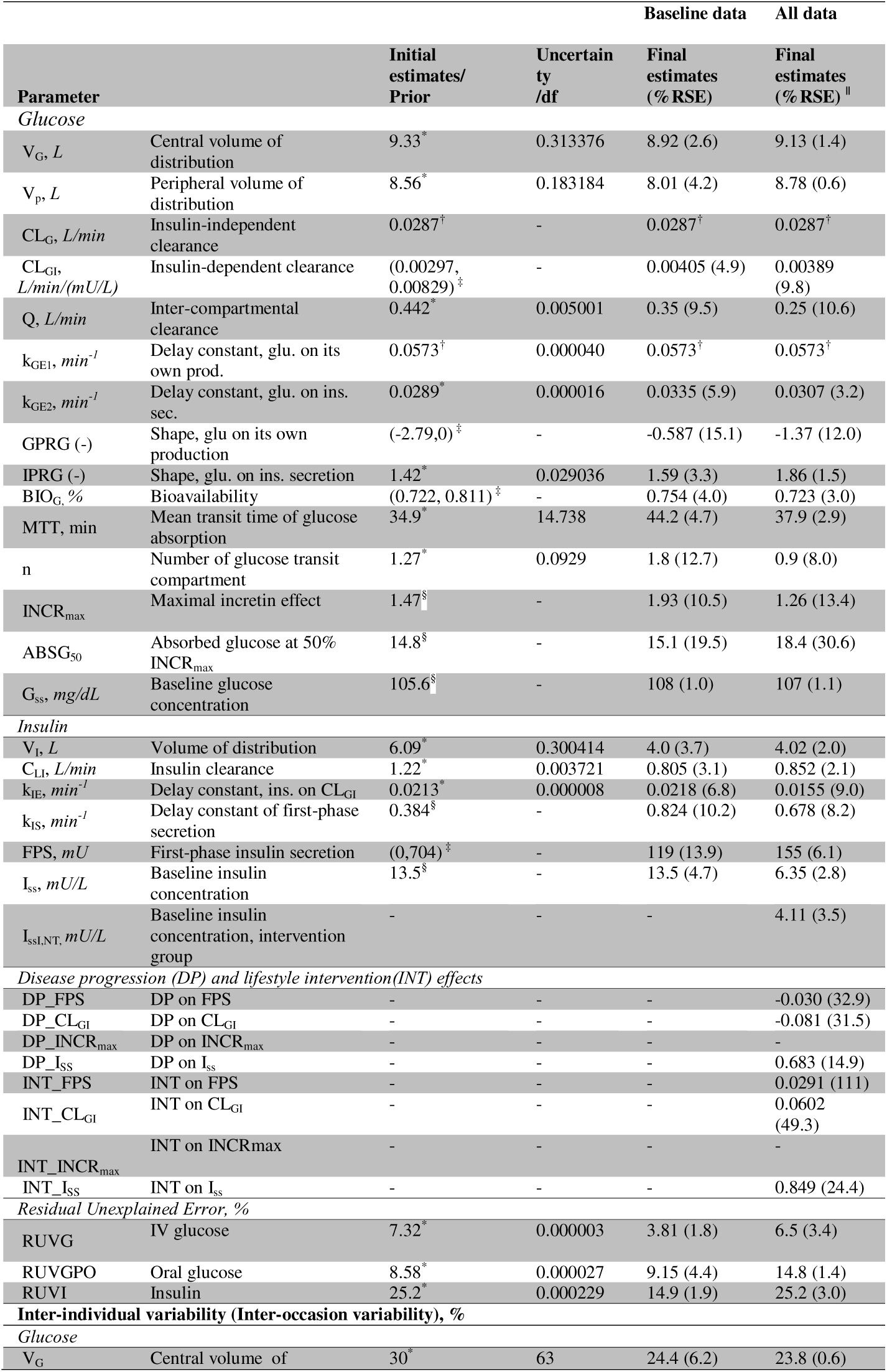
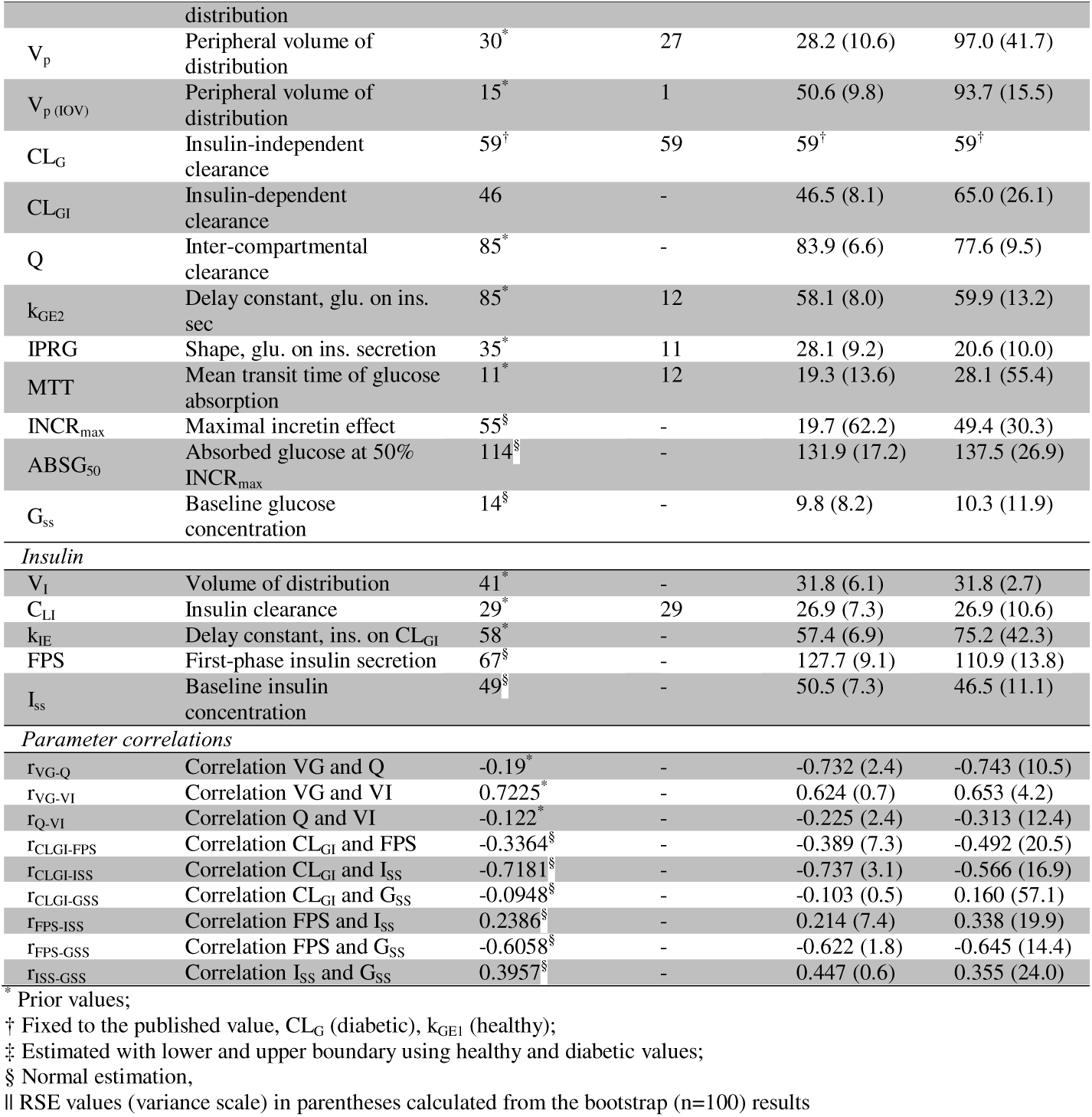
Final parameter estimates for only baseline and all data (baseline until the fourth year) in the combination of integrated glucose-insulin for intravenous and oral glucose tolerance test (IVGTT-OGTT-IGI) for the individuals with impaired glucose tolerance (IGT), using of prior information from oral and intravenous IGI model. ^1–3^

### The natural progression and lifestyle interventions

The data from baseline until the fourth year were used to model the natural DP and INT. DP model was set to initiate 24 hours after the end of Year 0 study in the dataset, allowing the glucose and insulin concentrations to return to the baseline values, before the next glucose provocations of the consecutive years. The DP and INT were investigated on the most reasonable parameters from a pathophysiological perspective. These parameters were the FPS as early insulin responsiveness in intravenous provocation, the CL_GI_ as a reflection of insulin sensitivity, I_SS_, reflecting changes in baseline insulin secretion and INCR_max_ as the changes of incretin hormones the diabetic state. The effect parameters were added one at a time (stepwise manner), to assess the significance of adding additional parameters in the nested models, using Likelihood Ratio Test (LRT) with the significance level of 5%.

Both step and linear models of DP (Equation 1a) and INT (Equation 1b) were tested, implemented according to Equation 1a and 1b. In the step models, changes were allowed between the baseline provocation at occasion=0 and the first occasion in the study, occasion=1, and thereafter no changes in the DP.

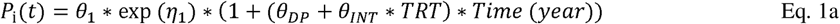

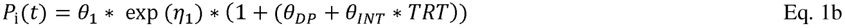

In both Equation 1a and 1b, θ_DP_ is the typical value of the natural disease progression for all individuals, without INT; θ_INT_ is the typical value of the lifestyle intervention effect. The parameter θ_INT_ is multiplied with an indicator variable, TRT; taking value 1 for the individuals in the intervention group, or 0 for the individuals in the control group. In Equation 1a, both θ_DP_ and θ_INT_ are interpreted as the slope of change with time, while in Equation 1b these parameters should be interpreted as the change between occasion 0 and 1. In the above equations, θ_1_ is a parameter with a progression with its log-normally distributed variability, η_1_, with a mean of 0 and variance of LJ ^2^.

### Specificity and sensitivity analyses

Sensitivity and specificity analyses were performed for the final model with DP and INT from the data of baseline until the fourth year, to assess the similarity between observed data and the model prediction in predicting who developed diabetes. The model-based diagnosis of diabetes was based on the diagnostic criteria of FPG or the 2-hour postprandial glucose values after a 75 g OGTT. There were 3 main scenarios investigated for the estimation process to produce the model prediction, which were:

A. using all the observed data of glucose and insulin concentrations,
B. using the data as in the scenario A, but excluding the glucose and insulin concentrations of the IVGTT at the fourth year, and
C. using the data as in the scenario A, but excluding the glucose and insulin concentrations at 0 and 120 minutes for OGTT.

For the calculation of the specificity and sensitivity analyses, the individuals who developed diabetes from the model prediction were compared to the ones who developed diabetes based on both the records of doctors’ diagnosis, and on the observed fasting and the 2-hour glucose concentrations from the data.

The sensitivity analysis represented the model ability to correctly identify the individuals who developed diabetes at the fourth year (Equation 2a). A value of 0.9 for example, showed the model’s ability to detect 90% individuals who developed diabetes (true positive) with 10% of individuals went undetected (false negative) by the model. Specificity analysis represented the model’s ability to correctly identify the individuals who did not develop diabetes until the fourth year (Equation 2b). For example, the value of 0.8 was translated to the ability of the model to identify 80% of the individuals who had not developed diabetes (true negative), with 20% falsely identified to develop diabetes (false positive), at the fourth year.^25^

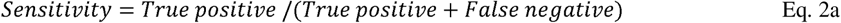

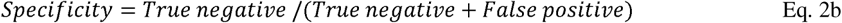

## Data analysis

Data analysis was done using nonlinear mixed-effects modeling with first-order conditional estimation with interaction (FOCE INTER) method in NONMEM (version 7.3)^23^ with ADVAN13 as the differential equation solver. Graphs were prepared and produced using the R software (version 3.3.1)^26^ as well as the ggplot2 package.^27^ A bootstrap run with 100 samples was used to assess the uncertainty in the parameter estimates. The visual predictive checks (VPCs)^28^ for different glucose provocations, randomization group and years were done as implemented in PsN (version 4.6.12) with 500 simulations, with the addition of censoring option for the individuals who develop diabetes, based on the FPG or the 2-hour plasma glucose concentration from the model prediction. This was done in order to mimic the original study design, in which the individuals who developed diabetes were excluded at the time of diagnosis.^19^ The visualization of the data and results was done using Xpose4 (version 4.5.0).^29–31^ The LRT was used to discriminate between nested models, with the chosen significance level of 5%. Based on LRT, the objective function value (OFV) is assumed to be chi-square distributed, and a decrease of 3.84 in OFV between hierarchical models with one parameter difference is considered statistically significant at 5% significance level. The final model was chosen based on the lowest OFV, the best VPCs for the full concentration-time profiles of glucose and insulin in each year, the baseline and the 2-hour glucose and insulin concentration-time profiles, scientific plausibility, as well as the best results from sensitivity and specificity analyses.

## Results

Final parameter estimates for the individuals with IGT were summarized in Table 1, under the baseline data final estimates. In general, the parameters with prior information were almost similar with or only slightly deviated from the initial value and for the parameters without prior information, the values were estimated to be in between the value of healthy individuals and patients with T2DM, with two exceptions. The rate of glucose effect on its own production (k_GE1_) was fixed to the previously published value of 0.0573 min^-1^ as it was observed to be unreasonably higher than the published value when estimated. The CL_G_ was fixed to the published diabetic value due its tendency to reach the lower boundary during estimation. The CL_GI_, FPS, and GPRG were closer to diabetic values, whereas BIO_G_ was closer to healthy values. The values for I_SS_ and G_SS_ agreed with the definition of IGT population. The results for every parameter were reasonable, translating to an adequate model description of the data, on the parameter level.

The inter-individual variability for some parameters as V_G_, INCR_max_, insulin clearance (CL_I_), peripheral volume of distribution (V_P_), effect delay rate constant of glucose on insulin secretion (k_GE2_), shape effect of glucose on insulin secretion (IPRG) and effect delay rate constant of insulin effect on glucose clearance (k_IE_) were decreased from the published values. The variability of the remaining parameters was either similar to or slightly higher than the published values. Parameter correlation between the inter-compartmental Q with V_G_ and insulin volume of distribution (V_I_) was increased in the IGT population, whereas the correlation between V_G_ and V_I_ was smaller.

To evaluate the model performance, the VPCs for the IVGTT-OGTT-IGI model in the IGT population are shown in Figure 2. The model fitted the median data of IVGTT and OGTT well for glucose and insulin component. An adequate model fit can be observed at the 2.5^th^ and 97.5^th^ percentile of the data for every component, especially for the insulin. However, by referring to the VPCs of glucose component in IVGTT, the variability of model prediction, which was at the 2.5^th^ and 97.5^th^ percentile, was a bit off from the observed data, especially at the end of time profile. This might be explained by a large heterogeneity in term of glucose control in this population, in which some of the individuals may be at the beginning of the IGT state close to the healthy state, whereas the other may be at the end, close to T2DM. Apart from that, this combination of IVGTT-OGTT-IGI model was able to describe the data of IGT population adequately.

**Figure 2.**
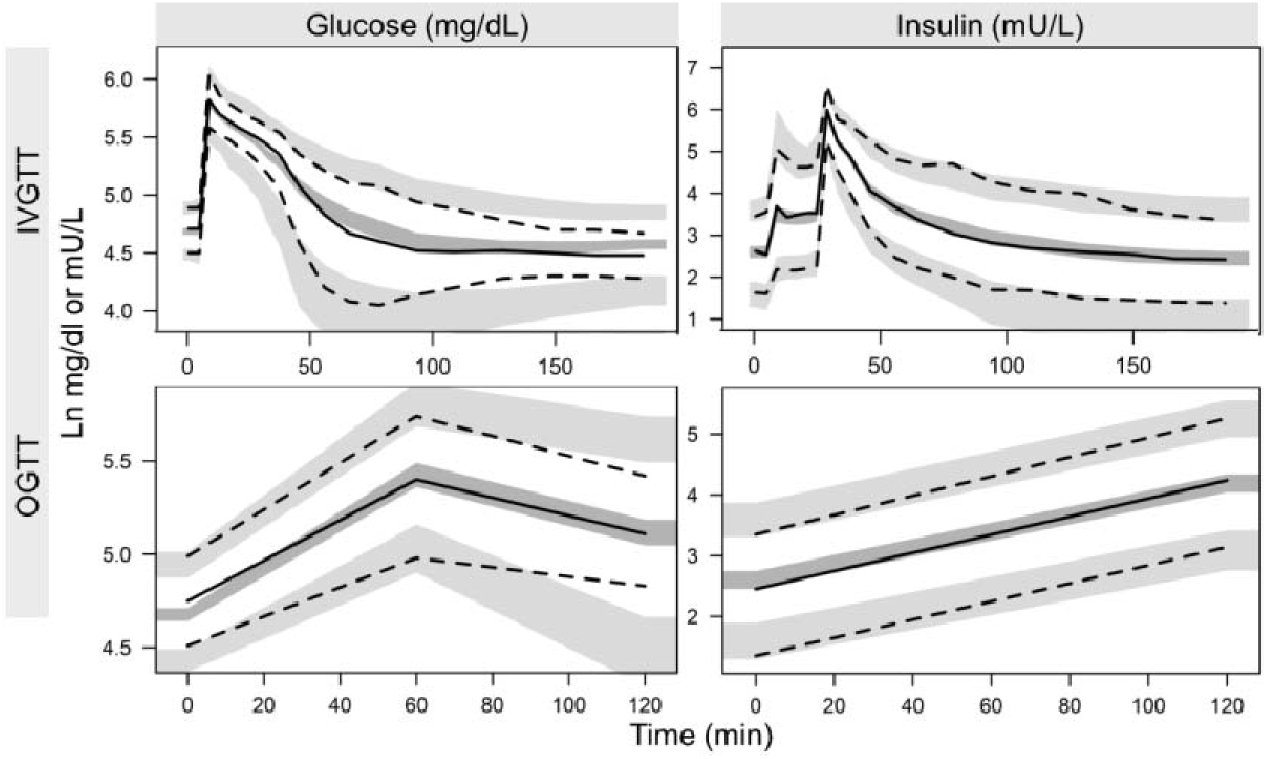
VPCs for the glucose and insulin for the individual with IGT, of the IVGG and OGTT. Solid lines represent the median of observed data, dashed lines represent the 2.5^th^ percentile (lower part) and 97.5^th^ percentile (upper part) for observed data. Dark shaded areas are the 95% confidence interval for median of simulated data, light shaded areas are the 95% confidence interval of 2.5^th^ percentile (lower part) and 97.5^th^ percentile (upper part) of the simulated data.

The effects of natural DP and INT on chosen parameters are summarized in Table 1, as are all final estimates. The FPS and CL_GI_ were decreased by 3.0% and 8.1% per year in all individuals because of the natural disease progression, whereas I_SS_ was increased 68.3% from baseline to Year 1, and remained unchanged after the first year. On top of the natural disease progression, the lifestyle intervention affected the FPS value by 2.9% increase per year, producing a net effect of 0.1% decrease per year. This indicates a much slower deterioration of the acute response of beta-cell function to produce insulin, following an intravenous glucose load. For the CL_GI_, an increase by 6.0% per year with lifestyle intervention was observed, producing a 2.1% decrease per year, representing a slower deterioration of insulin sensitivity. The I_SS_ was affected by the INT with a net effect of 153% increase from baseline to the first year and remained constant after that, reflecting the improvement of beta-cell function to increase insulin secretion at that time.

For the investigations on the INCR_max_, no significant effects were found. The model fit with DP on incretin was no better than the chosen final model, in which an insignificant OFV improvement and worse visual predictive checks were observed. Besides, the specificity and sensitivity results were also worse for the model with DP on incretin than the chosen final model.

The VPCs for glucose and insulin component in control and intervention groups are shown in Figure 3 for IVGTT.

**Figure 3.**
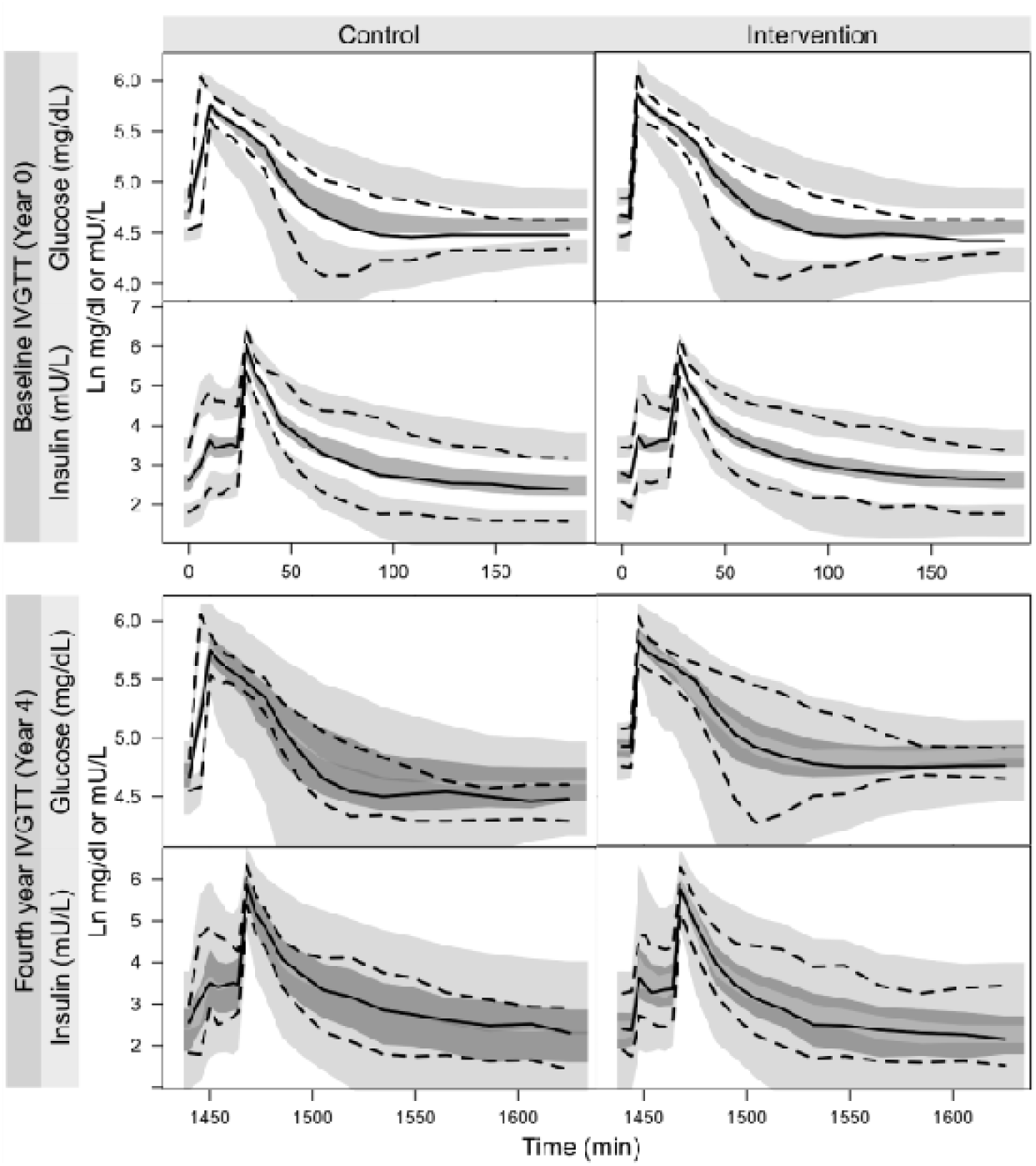
VPCs for frequently sampled IVGTT from baseline and the fourth year for final model of IGI among individuals with IGT, with the diabetes DP and INT. Solid lines represent the median of observed data, dashed lines represent the 5^th^ percentile (lower part) and 95^th^ percentile (upper part) for observed data. Dark shaded areas are the 95% confidence interval for median of simulated data, light shaded areas are the 95% confidence interval of 5^th^ percentile (lower part) and 95^th^ percentile (upper part) of the simulated data.

Based on the figure, the IGI model, including natural DP and INT, has well described the data for the baseline and the fourth year for IVGTT. Overall model fit was adequate for both glucose and insulin components at baseline and in the fourth year, among individuals in both the control and lifestyle intervention groups. The median and the 5^th^ percentile model prediction were very good for all cases. Only a small misfit was observed in the 95^th^ percentile of the model prediction for the baseline glucose component, where a model over-prediction is observed. A possible explanation might be due to the high variation of glucose concentrations in the population, related to whether individuals were close to healthy or diabetic status. On the fourth year, the confidence interval for the model prediction was wider than baseline, as the individuals who developed diabetes were excluded from the VPC analysis. The VPCs for OGTT are provided as a supplementary material (Supplement S1). A very good model fit can be observed on the median, 5^th^ and 95^th^ percentile data in all cases, except for the 5^th^ percentile of G_ss_ component, which was over-predicted by the model. The same possible explanation as for the misfit of baseline glucose for IVGTT may be applied in this case.

Besides than being able to describe the full glucose and insulin concentration-time profiles, the final model was able to adequately describe the baseline and the 2-hour glucose and insulin, as illustrated in Figure 4.

**Figure 4.**
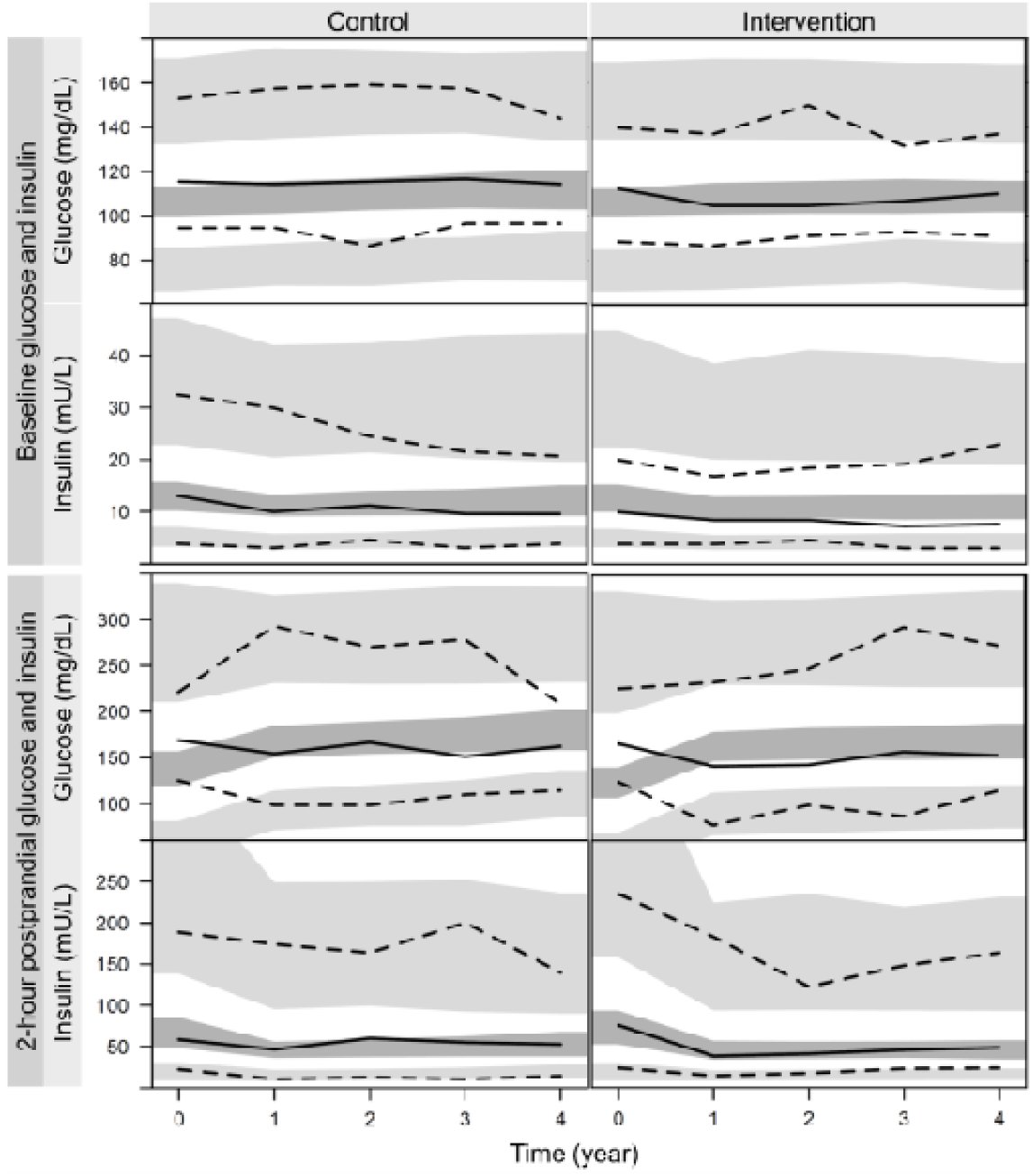
VPCs for baseline and 2-hour plasma glucose and serum insulin concentration for final model of IGI among individuals with IGT, with the diabetes DP and INT. Solid lines represent the median of observed data, dashed lines represent the 2.5^th^ percentile (lower part) and 97.5^th^ percentile (upper part) for observed data. Dark shaded areas are the 95% confidence interval for median of simulated data, light shaded areas are the 95% confidence interval of 2.5^th^ percentile (lower part) and 97.5^th^ percentile (upper part) of the simulated data.

Specifically, the model fit for the 2.5^th^ percentile of the baseline glucose was a bit underpredicted in the control group. For the 2-hour glucose concentration, a misfit can be found at baseline (Year 0), in which the glucose was underpredicted. This misfit may be explained by a condition known as the regression toward the mean.^32^ The individuals were selected based on their 2–hour postprandial glucose in between 160-220 mg/dL, based on the IGT criteria from the WHO technical report series 1985^21^, therefore, a tight variability for the baseline was observed with a specific mean value. However, for the subsequent observations (for example at the first year), the variability may increase by chance, in which the individuals who had a value close to the mean at the first occasion, had a value that was further from the mean at the second one. For the insulin component, a small misfit can be found at the 97.5^th^ percentile and the later time point for the median data.

In terms of the performance of the model prediction, the results of sensitivity and specificity are summarized in Table 2, incorporating the analyses calculated based on the doctors’ diagnosis as well as the observed data. Based on the doctors’ diagnosis, the sensitivity analysis result was the best for scenario B (86%), which was by excluding the glucose and insulin concentrations for the IVGTT at the fourth year, followed by scenario B, by using all data (82%) and finally scenario C, by excluding the glucose and insulin concentrations at 0 and 120-minutes (68%). Even though scenario B had the best sensitivity, it had a lower specificity (87%) than scenario A (95%) but slightly higher than scenario C (84%). For the analyses results based on the observed data, a much lower sensitivity but a small difference in the specificity results as compared to the doctors’ diagnosis for all scenarios were obtained. About 47% sensitivity was recorded when using all data (scenario A) and excluding the glucose and insulin concentrations at the fourth year (scenario B), and 36% when excluding the glucose and insulin concentrations at 0 and 120 minutes (scenario C). However, for the specificity, higher values were generally obtained for scenario A, B and C, which were 91%, 87% and 78%, respectively.

**Table 2.** The summary of the sensitivity and specificity analyses result based on the diabetes diagnosis made by the doctors’ (Doctors’ diagnosis) and by referring to the observed fasting and 2-hour glucose concentration (Observed data), in 3 different scenarios: A, B, and C.

| Scenario | Sensitivity |  | Specificity |  |
| --- | --- | --- | --- | --- |
|  | Doctors' diagnosis | Observed data | Doctor's diagnosis | Observed Data |
| Scenario A: Using all glucose and insulin concentrations | 0.82 | 0.47 | 0.95 | 0.91 |
| Scenario B: Excluding the glucose and insulin concentrations for IVGTT at the fourth year | 0.86 | 0.47 | 0.87 | 0.87 |
| Scenario C: Excluding the glucose and insulin concentrations at 0 and 120-minutes | 0.68 | 0.36 | 0.84 | 0.78 |

## Discussion

The IGI model has previously been published with distinct estimates for OGTT and IVGTT for diabetic and healthy populations.^1–3^ In the current study with an IGT population, NONMEM’s prior functionality was considered the best approach for determining the model parameters regardless of the type of glucose provocation or health status, as the functionality incorporates the published parameter values and their uncertainties during estimation. The final parameter estimates for the IGT population are comparable to the published values. Besides, this approach has been shown to produce a more stable analysis, especially with complex models as the IGI model. ^22^

CL_G_ approached the lower boundary during estimation, i.e., the diabetic value. This could indicate similarity with patients, but it could also be due to its non-identifiability in the absence of labelled glucose data. Labelled glucose was available during the development of the first IGI model,^1–3^ which supports differentiation between provocation glucose and endogenous glucose. Related to the absence of labelled glucose, the effect of glucose on its own production was difficult to determine; the rate, k_GE1_, was estimated to be immediate, and the shape, GPRG, was close to zero. Notably, for the T2DM population, the effect of glucose on its own production is absent. In summary, on the population level, the insulin sensitivity (i.e. CL_GI_), insulin secretion function following a high glucose load (i.e., FPS), and feedback mechanism function of glucose on its own production (GPRG) were close to the state of diabetes mellitus. However, the glucose bioavailability (BIO_G_) was similar to the healthy state.

Over time, FPS, CL_GI_, and steady-state insulin (i.e., I_SS_) changed in the IGT population; however, differently for the control and the intervention groups. The FPS decreased each year, indicating deterioration in acute beta-cell function’s response to secrete insulin following an intravenous provocation with a high glucose concentration. Similar results were obtained in previous studies, which reported significant reduction of FPS in the IGT individuals when using IVGTT^6,33–35^ and hyperglycemic glucose clamps.^10,36,37^ Additionally, a study concluded a higher reduction of the FPS in the IGT population with first-degree relatives with T2DM than those without heredity.^38^ Different studies reported different reduction rates of the FPS, most probably due to differences in the type of glucose provocations, genetic factors, and tissue sensitivity to insulin.^33^ CL_GI_ decreased over time due to the natural disease progression, indicating a reduction of the insulin sensitivity over the years. This was in agreement with other studies’ results that reported a reduction in the insulin sensitivity translated from 20-30% decrease of the CL_GI_ among individuals with IGT, compared to individuals with NGT,^6,34,35,39,40^ even after BMI adjustment^34^. The combination of insulin secretion depletion and increased muscle insulin resistance was determined to be the most prominent cause of decreasing insulin sensitivity in the IGT population.^33^ Moreover, the I_SS_ increased from the baseline to the first year. This may be a compensatory mechanism of beta-cells to secrete more insulin as response to the required higher need with decreasing CL_GI_ (i.e., increasing insulin resistance), especially in the early progression of T2DM. In this condition, more insulin is needed to suppress the increase of baseline glucose concentration. This feedback mechanism phenomenon has been well documented.^41^ It was also reported in previous studies that fasting plasma insulin concentration was higher in the IGT than in NGT population.^6,33,34,36,42^ Another study revealed a higher baseline insulin secretion rate among the IGT, than the NGT counterpart.^37^

On top of the natural disease progression, the intensive lifestyle intervention with counselling on diet, weight reduction, and exercise were quantified to influence the physiological changes of IGT individuals. The acute insulin secretion from pancreatic beta-cells had a much slower deterioration, marked by almost no change in FPS. In addition, a slower deterioration of insulin sensitivity was observed with intervention, translated from the final net effect of a smaller reduction of the parameter value with the intervention, over time. Apart from the acute insulin response and insulin sensitivity, the beta-cell function in secreting basal insulin was further improved by the intervention. This was indicated by a net incremental basal insulin level from baseline until the first year. The possible explanation might be related to the physiological improvement in beta-cell function to secrete insulin at baseline, with the effect of the lifestyle intervention. These results complement those from previous studies on the benefits of lifestyle interventions (dietary counselling, weight reduction, and moderate-intensity physical activity for at least 150 minutes per week) on insulin sensitivity and insulin secretion, resulting from improved beta-cell function, among individuals with IGT.^15–17^ A study reported the improvements of glucose tolerance and insulin sensitivity in the lifestyle intervention group, determined by the weight loss and increased physical fitness.^15^ Another study concluded that the lifestyle intervention significantly reduced the incidence of diabetes among individuals with IGT, due to improved beta-cell function and insulin sensitivity.^16^ In addition, a 58% reduction in progression from IGT to T2DM during a mean follow-up of 2.6 to 3.2 years with the effect of lifestyle intervention was documented in two studies.^18,19^

The final model, without inter-individual variability, for the natural progression and the intervention best described the data. This result is not in agreement with current beliefs; the rate of progression to diabetes and the extent of interventions are different between individuals in relation to the individuals’ insulin sensitivity and beta-cell function, weight and genetic factors.^16,17,33^ The interindividual variability in the IGI model, at baseline, is fairly large and the inability to estimate additional interindividual variability related to the progression over time might be related to the flexibility of the model without disease progression. Disappointingly, the inclusion of markers for intervention success, e.g., extent of weight loss, or adherence to intervention recommendations did not improve the model fit (results not shown).

In theory, the incretin effect should also be affected by the natural progression and intervention in this population. Incretin potentiation of glucose-mediated insulin secretion is known to deteriorate as diabetes progress over time, which may be related to a progressive weakening the ability of incretin hormone in the form of glucose-dependent insulinotropic peptide (GIP) and glucagon-like peptide-1 (GLP-1), during the progression of T2DM, to stimulate glucose-dependent insulin secretion after oral glucose intakes.^43–47^ In some studies, the GIP was reported to have entirely lost its ability to modulate the glucose-dependent insulin secretion in the patients with T2DM, whereas GLP-1 still exhibits its insulinotropic function in the patients’ population, although with an efficiency lower than healthy individuals.^43–46,48^ Another study documented a 25% impairment of GLP-1 response to an OGTT in prediabetes as compared to individuals with NGT.^12^ The lifestyle interventions were suggested to improve the incretin effect, for example as an effect of changes of insulin secretion directly related to the GIP response to an oral glucose tolerance test.^49^ However, modelling of natural progression and intervention on incretin function did not show significantly changes in the current study. This might be due to an insufficient time frame in the dataset for the model to capture the changes that usually developed after 5 to 10 years.^50,51^ Additionally, changes in incretin effects have been reported for patients who already have diabetes and the deterioration of incretin effect may only occur at a more advanced state of the disease. Other possible reasons include insufficient OGTT data to quantify those changes, as well as the complexity of the IGI model, which may result in identifiability issues. As only sparse sampling was performed in the OGTT, the relative importance of fitting OGTT data in relation to IVGTT data may be reduced. To support estimates of incretin effects, a study design in which the OGTT is matched with an isoglycemic intravenous glucose test would have provided substantial information to quantify changes over time. In the current study, a linear time-varying FPS was found to produce the best model fit based on the model diagnostics criteria. As no IVGTT data, with FPS, was available for the first until third year, the estimation of a linear relationship may be questioned. Initially, a step function was implemented, with the assumption of unchanged FPS from baseline until the third year and later increased at the fourth year. However, the implementation failed to produce good results, with for example a massive over-prediction of 2-hour glucose concentration in the VPCs, and scientifically implausible parameter estimations. Therefore, the linear DP to describe the FPS changes was considered rational, supported by the physiological reasons as discussed previously in the current study.

The sensitivity and specificity analyses were conducted across 3 scenarios: A) using all data, B) excluding data from the IVGTT at the 4^th^ year, and C) excluding glucose and insulin concentrations at 0 and 2 hours. Scenario A represents a best-case scenario of the sensitivity and specificity analysis, as all data were used in the estimation process. Scenario B was investigated to remove the influence of the IVGTT data at the fourth year. Scenario C was designed to exclude data that are used to diagnose diabetes (glucose at time 0, i.e., fasting glucose) and impaired glucose tolerance (2 h glucose), and let the model predict the missing observations, based on the model. Thus, in scenario C, predictions of measurements for diagnosis were purely model-driven. Overall, results showed acceptable sensitivity and specificity compared with the diagnostic criteria as specified in the study. The final model was able to identify 68% to 86% individuals who developed diabetes, and only 14% to 32% went undetected (sensitivity analysis). The model was also able to identify 84% to 95% individuals who did not develop diabetes, with only 16% to 5% falsely identified to develop diabetes at the 4^th^ year (specificity analysis). These results suggest that the glucose and insulin concentrations at the 4^th^ year may have an influence on the specificity, but not sensitivity analysis, by referring to the 86% sensitivity and 87% specificity for the scenario B, as compared to 82% sensitivity and 95% specificity for the scenario A. Besides, the model was able to predict acceptable 0 and 2 h glucose and insulin concentrations from the baseline until the 4^th^ year, given the high sensitivity (68%) and specificity (78%) of the model, in scenario C.

Disappointingly, the sensitivity and specificity analyses results based on the actual diagnoses showed quite a poor sensitivity with less than 50% of the sensitivity results obtained for all scenarios. The specificity, however, was good, ranging from 78% to 91%, indicating a high model performance in correctly identifying individuals who never progressed to diabetes, but quite poor at identifying those who developed diabetes at the 4^th^ year. It was noted that if the diagnostic criteria for diabetes were used based on the data, more individuals would have been deemed diabetic, and that the physicians did not use the criteria strictly. This may be related to the fact that a second OGTT was performed to confirm the diabetes diagnosis, if any individuals had high fasting or 2-hour glucose levels. The data from the second OGTT were unfortunately not available for this analysis. Most probably in a high number of cases, the second OGTT ruled out a diabetes diagnosis; therefore, fewer individuals were diagnosed diabetic, as compared to when the diagnosis was made directly from the observed data.

The natural progression and intervention model developed in this study can be used to quantify the effects of non-medication intensive lifestyle intervention on the changes on the glucose and insulin regulations among the IGT population, using the IGI model. Further investigations on the effects of the medication intervention would be worth doing.

## Conclusion

A natural DP model was successfully added in the combination of IVGTT and OGTT IGI model to describe parameter changes of FPS, CL_GI_, and I_ss_ among the population with IGT, with the quantification of the effects of lifestyle intervention. In particular, the FPS and CL_GI_ were deteriorated in a much slower rate after the intensive lifestyle intervention.

## Acknowledgements

The research leading to these results has received support from the Innovative Medicines Initiative Joint Undertaking under grant agreement n° 115156, resources of which are composed of financial contributions from the European Union’s Seventh Framework Programme (FP7/2007-2013) and EFPIA companies’ in kind contribution. The DDMoRe project is also financially supported by contributions from Academic and SME partners.

