## Supplementary material S1 for "Mathematical diabetes disease progression modeling in the integrated glucose-insulin model among individuals with impaired glucose tolerance from the Finnish Diabetes Prevention Study"

### Supplement S1

VPCs for OGTT from baseline until the fourth year for final model of integrated glucose-insulin (IGI) among individuals with IGT, with the diabetes DP and INT. Solid lines represent the median of observed data, dashed lines represent the 5^th^ percentile (lower part) and 95^th^ percentile (upper part) for observed data. Dark shaded areas are the 95% confidence interval for median of simulated data, light shaded areas are the 95% confidence interval of 5^th^ percentile (lower part) and 95^th^ percentile (upper part) of the simulated data.

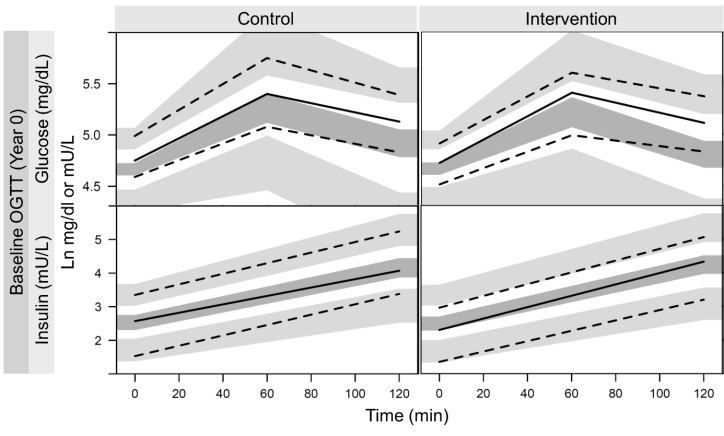

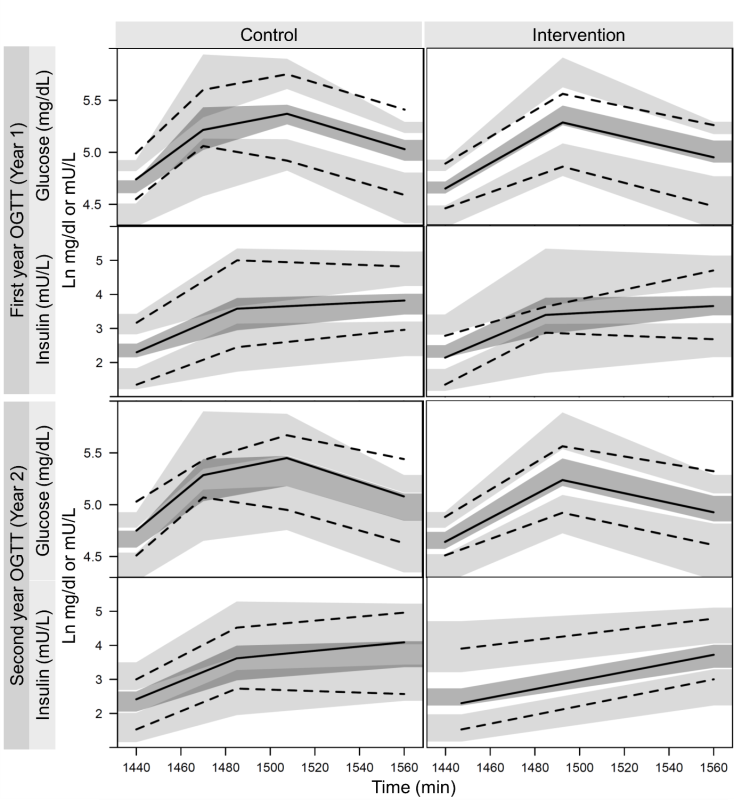

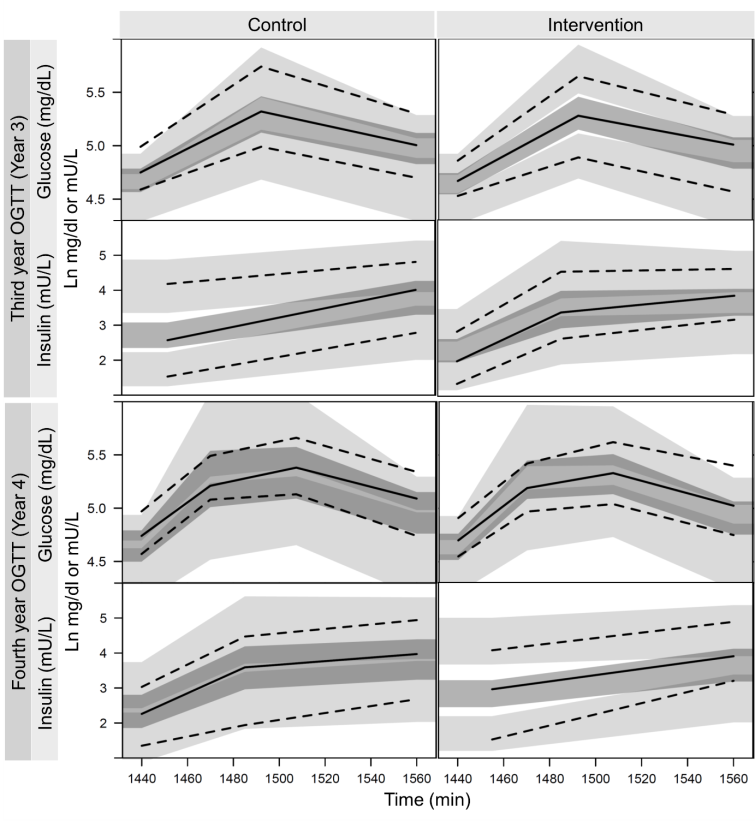

**Supplement S1.** VPCs for OGTT from baseline until the fourth year for final model of integrated glucose-insulin (IGI) among the individuals with IGT, with the diabetes disease progression and lifestyle intervention effects. Solid lines represent the median of observed data, dashed lines represent the 5^th^ percentile (lower part) and 95^th^ percentile (upper part) for observed data. Dark shaded areas are the 95% confidence interval for median of simulated data, light shaded areas are the 95% confidence interval of 5^th^ percentile (lower part) and 95^th^ percentile (upper part) of the simulated data.
